# Establishment of a novel set of vectors for transformation of the dinoflagellate *Amphidinium carterae*

**DOI:** 10.1101/2022.12.15.520308

**Authors:** I.C. Nimmo, K. Geisler, A.C. Barbrook, F.H. Kleiner, A. Scarampi, D. Kosmützky, R.E.R. Nisbet, C.J. Howe

**Affiliations:** Department of Biochemistry, University of Cambridge, Tennis Court Road, Cambridge, CB2 1QW, UK; Cambridge Institute of Therapeutic Immunology and Infectious Disease, University of Cambridge, Jeffrey Cheah Biomedical Centre, Cambridge Biomedical Campus, Puddicombe Way, Cambridge, CB2 0AW, UK; Department of Plant Sciences, University of Cambridge, Downing Street, Cambridge, CB2 3EA, UK; School of Biosciences, University of Nottingham, Sutton Bonington Campus, Sutton Bonington, Loughborough, Leics. LE12 5RD, UK; Department of Bionanoscience, Delft University of Technology, Gebouwnummer 58, Van der Maasweg 9, 2629 HZ, Delft, The Netherlands

## Abstract

Peridinin-containing dinoflagellate algae have a chloroplast genome formed from plasmid-like minicircles. This fragmented genome has allowed us to develop a genetic modification methodology involving the use of biolistics to introduce artificial minicircles in *Amphidinium carterae* (Nimmo et al., 2019). The previously reported artificial minicircles were based on native minicircles containing either the *psbA* or *atpB* gene. Each artificial minicircle allowed expression of a single selectable marker instead of *psbA* or *atpB*. Here, we present two further artificial minicircles for use in transformation of *A. carterae*. One is based on the *petD* minicircle, allowing the expression of a single selectable marker. The second is based on the two-gene minicircle originally containing *atpA* and *petB*, and allows the dual expression of a selectable marker and a gene of interest. Our research suggest that all of the 20 or so minicircles in *A. carterae* are suitable for adaptation as artificial minicircles, allowing for the simultaneous introduction of multiple genes.

## Introduction

The ability to alter genetic material has revolutionized cell biology. The majority of modern biochemical analyses are carried out in cells which have been genetically modified to introduce, remove or mutate genes. However, these analyses rely on the establishment of a reliable genetic modification protocol. Until recently, it was not possible to modify genetically one of the largest and ecologically most important groups of eukaryotic groups micro-organisms, the dinoflagellate algae (Chen et al., 2019). Dinoflagellate algae are found in aquatic environments worldwide, and are an essential symbiont in coral reefs.

In 2019, we established the first genetic modification system in dinoflagellates, in *Amphidinium carterae* (Nimmo et al., 2019). *A. carterae* has been used as a model organism for dinoflagellates, as it lacks the armoured thecal plates common in other species. These thecal plates have been proposed to be one reason why genetic transformation of dinoflagellates has proved difficult (Chen et al., 2019). To establish transformation, we made use of the unique fragmented dinoflagellate chloroplast genome. Instead of having a conventional 120-150 kb circular chloroplast genome, many dinoflagellate species have a fragmented chloroplast genome of approximately 20 plasmid-like minicircles (Howe et al., 2008). Dependent on species, each minicircle contains between 0-4 genes and is up to 5 kb in size. Each contains a conserved core region containing the origin of replication and transcriptional start site (Howe et al., 2008). By replacing the protein-coding gene in the *psbA* or *atpB* minicircles with a selectable marker, we were able to create artificial minicircles (as shuttle vectors) for genetic transformation (Faktorová et al., 2019; Nimmo et al., 2019).

Here, we present a further two artificial minicircles, suitable for introducing genes into the dinoflagellate chloroplast. One is based on the *petD* minicircle, and the other is based on the miniciricle containing *petB* and *atpA* genes. The latter allows for the expression of two transgenes. These results are a significant advance in the toolset available for dinoflagellate transformation, and will allow researchers to answer important biological questions.

## Materials and Methods

### Growth of A. carterae

Cultures were maintained as described in Nimmo et al., 2019.

### Creation of constructs

All constructs were based on the native *petD* and the *petB/atpA A. carterae* minicircles (Genbank accessions AJ250265 and AY048664), as shown in Figure 1. For the construct pAmpPetDChl, the *petD* gene was replaced with a sequence encoding chloramphenicol acetyl transferase (CAT; codon optimized for *A. carterae*), and lineariezed after the end of CAT. For the two-gene construct, the *petB* gene was replaced with CAT, and the *atpA* gene was replaced with a sequence encoding green fluorescent protein (eGFP, condon optimized for *A. carterae*) to give the construct pAmp2GChlGFP. Both sequences were synthesized by GeneArt (Invitrogen) and inserted into the *E. coli* pMA cloning vector. All construct sequences are given in Suppl. data.

**Figure 1.**
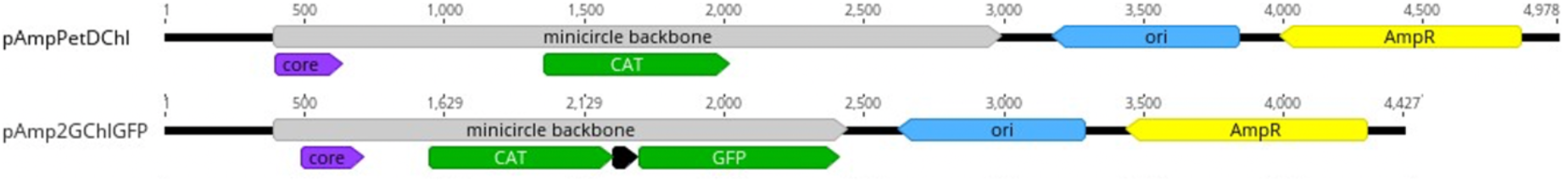
Linearized artificial minicircles pAmpPetDChl and pAmp2GChlGFP. The grey arrow indicates region of minicircle origin, the remainder is bacterial shuttle vector (based on the *E. coli* vector pMA). The bacterial origin of replication (*ori*) is blue and the minicircle core region (core) is purple. AmpR (yellow) is ampicillin resistance for selection in *E. coli* and CAT (green) is chloramphenicol acetyl transferase, for selection in *A. carterae*. GFP (green) is eGreen Fluorescent Protein, codon optimized for *A. carterae*. The intergenic region (black) is the original intergenic region from the *A. carterae* two-gene minicircle encoding *petB* and *atpA*.

### Transformation

Biolistic transformation was carried out as developed by Nimmo et al, 2019, using 1550psi rupture disks, and chloramphenicol selection at 20µg/ml (in DMSO) was applied from three days following particle bombardment. A full protocol can be found at https://www.protocols.io/view/biolistic-transformation-of-amphidinium-4r2gv8e

### PCR and RT-PCR

Extraction of DNA and RNA and PCR and RT-PCR (including primer sequences) are described in Nimmo et al, 2019. Primers for detection of GFP:

Fwd- GGTGATGTAAACGGTCATAAG; Rev- GTGTATCACCCTCGAACTTTAC, giving rise to a 298 bp product. Primers for pAmpPetDChl:

Fwd- CGACAGGAGTGATTAACTGAG;

Rev- GGAATGCGGTGATATCAAGCTGTACTG, giving rise to a 1061bp product. Primers for pAmp2GChlGFP: Fwd - GGAATGCGGTGATATCAAGCTGTACTG

Rev- CGGAACTCTGGATGTGCG giving rise to a 430bp product.

## Results

### Development of new transformation vectors

We designed two new artificial minicircles as transformation vectors, pAmpPetDChl and pAmp2GChlGFP. The pAmpPetDChl vector is based on the *petD* single gene minicircle, while the pAmp2GChlGFP vector is based on the two-gene minicircle containing *petB* and *atpA*. Each artificial minicircle has an *E. coli* pMA plasmid backbone, and contain the full minicircle sequence, with either the *petD* or *petB* coding sequences replaced with a sequence encoding chloramphenicol acetyl transferase (CAT). In the two-gene minicircle, the *atpA* coding sequence is replaced with one for eGFP. These vectors are designed to allow selection in *E. coli* with ampicillin, and for selection in *A. carterae* with chloramphenicol. Note that selection in *E. coli* with chloramphenicol was not successful.

To determine if there was a difference in transformation success rates between artificial minicircles, *A. carterae* cells were bombarded with each artificial minicircle multiple times, followed by selection with chloramphenicol, as shown in Table 1. PCR to test for presence of each artificial minicircle was carried out following each transformation (Figure 2). We included our two previously published minicircles pAmpChl (based on the *A. carterae psbA* minicircle (Nimmo et al., 2019)) and pAmpAtpBChl (based on the *A. carterae atpB* minicircle (Faktorová et al., 2019)) for comparison. Although the success rates with which transformed cells were recovered varied from 33% to 75%, there was no statistical difference between vectors (Chi-squared, p=0.05), as shown in Table 1.

**Table 1.** Efficiency of transformation. Each artificial minicircle was used to transform wild type *A. carterae* in separate biolistic transformations. The number of times that cultures contained transformed cells (as confirmed by PCR) was recorded. Each transformation experiment (or set of experiments) also included a control, to ensure that cells survived the biolistic procedure. Transformations were carried out at both the University of Cambridge and University of Nottingham by separate individuals, and transformants were obtained at both sites.

| Artificial minicircles | Original minicircle | Size (bp) | Times shot | Successful transformation | % success |
| --- | --- | --- | --- | --- | --- |
| pAmpChl | *psbA* | 4326 | 12 | 9 | 75% |
| pAmpAtpBChl | *atpB* | 3870 | 9 | 4 | 44% |
| pAmpPetDChl | *petD* | 4978 | 9 | 3 | 33% |
| pAmp2GChlGFP | *atpA/petB* | 4427 | 13 | 5 | 39% |

**Figure 2.**
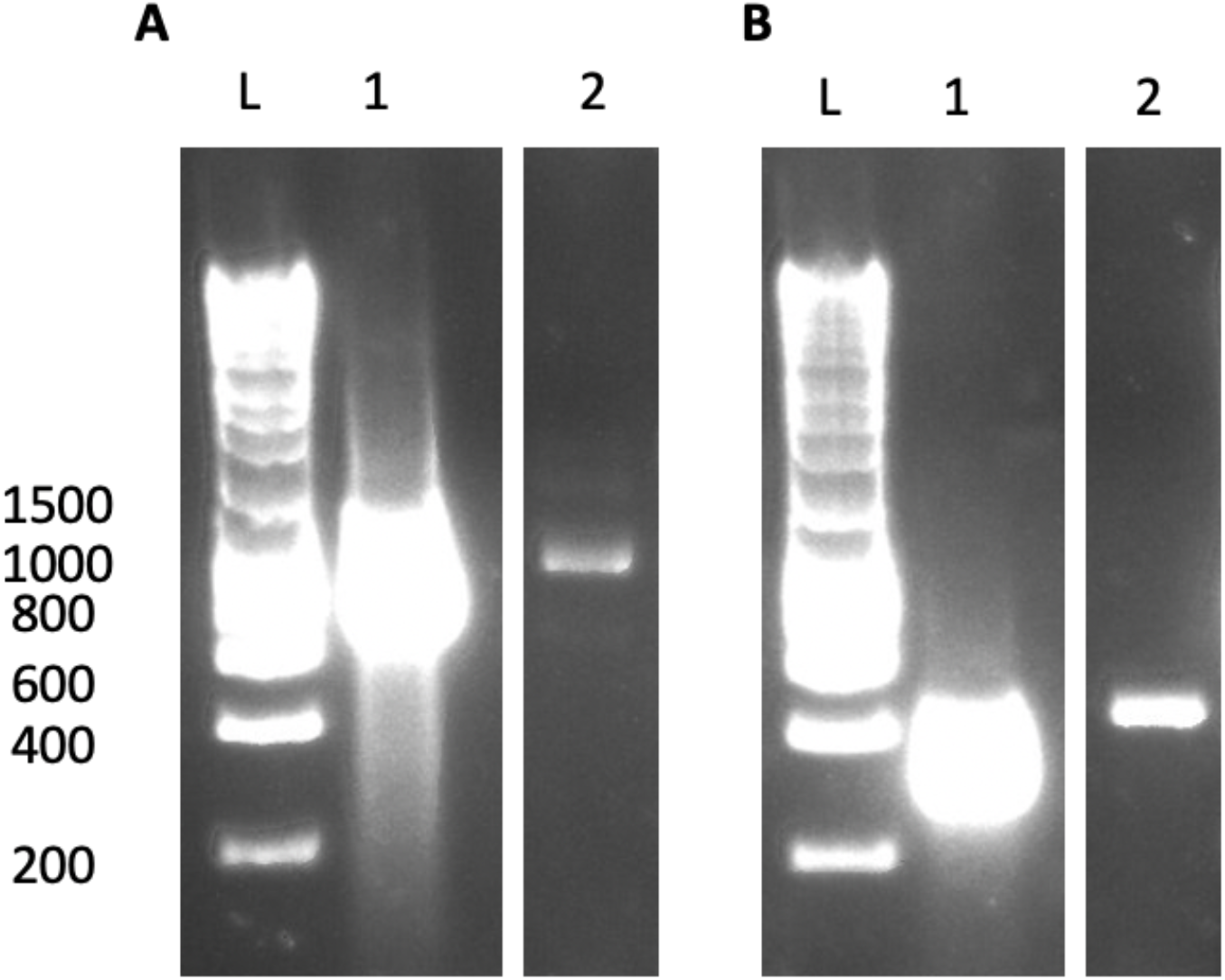
PCR to test for successful transformation. **A.** pAmpPetDChl, **B.** pAmp2GChlGFP Lane L, hyperladder 1kB (Bioline), Lane 1 positive control (purified transformation vector), Lane 2, DNA isolated from putative transformed *A. carterae*.

### Effect of artificial minicircle on A. carterae growth rate

To determine how the introduction of artificial minicircles affected the growth rate of *A. carterae*, cultures of each of the four lines were established in triplicate, and grown in standard conditions in the presence of chloramphenicol for 21 days. This showed that the strains containing pAmpChl and pAmpAtpB had similar growth rates, while strains containing the pAmpPetDChl and pAmp2GChlGFP grew at substantially lower rates, Figure 3.

**Figure 3.**
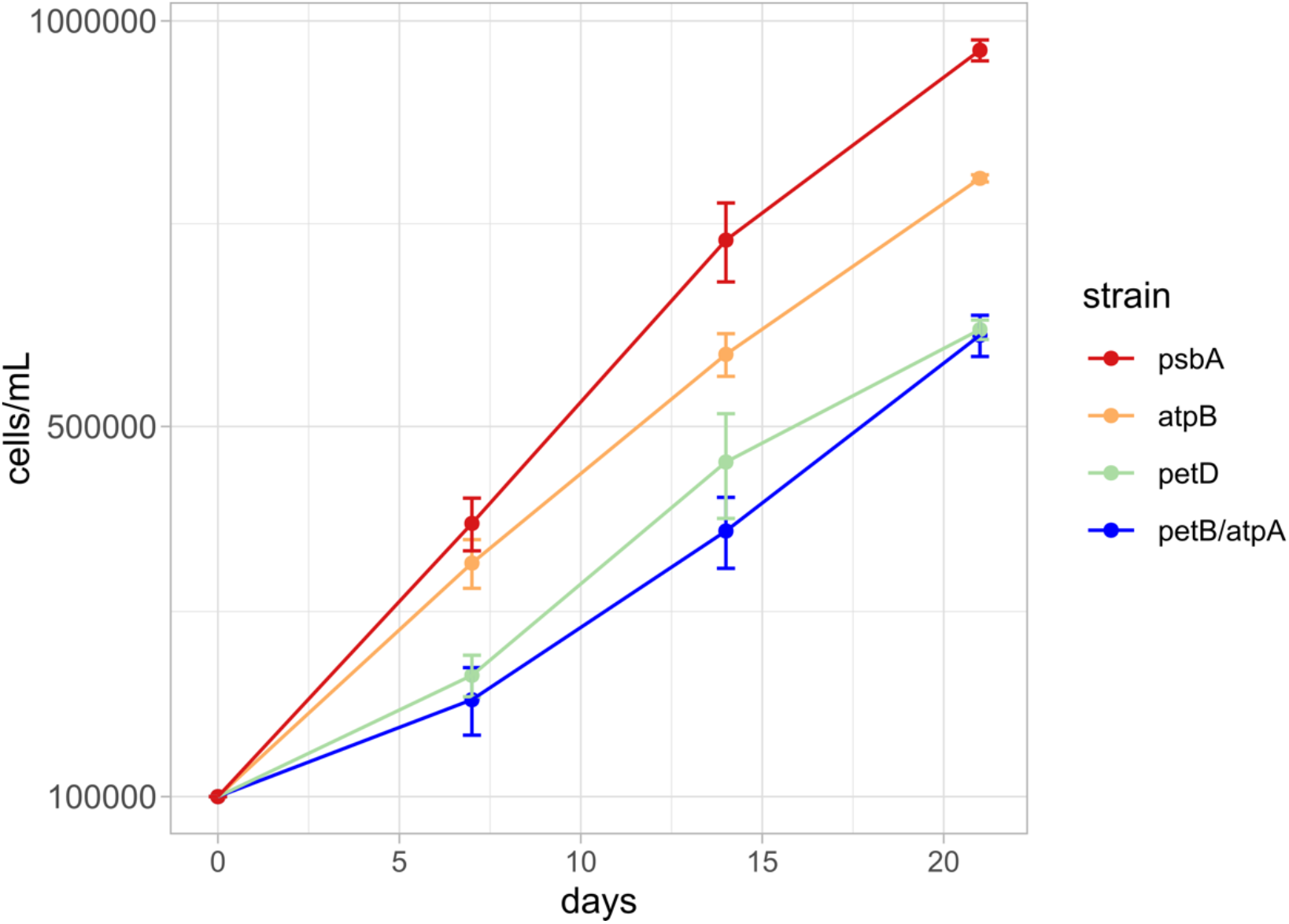
Growth curves. The four transformed cell lines were grown (in triplicate) in the presence of chloramphenicol for 21 days. Cell counts were undertaken using a haemocytometer at days 7, 14 and 21. The original minicircle on which each construct was based is shown. All cultures contained 100,000 cells at day 0.

We tested if average growth rates of cells over 21 days with the chloramphenicol resistance gene replacing different endogenous genes correlated with the transcript levels of the endogenous genes in wild type cells. The growth rate of cultures resulting from transformation with each artificial minicircle was compared with known transcript levels of each native minicircle, as shown in Table 2. The transcript level data were obtained from the same strain in our laboratory (Cambridge) in previous experiments (Dorrell et al., 2019). The results showed that cultures transformed with plasmids based on minicircles corresponding to more abundant transcripts (pAmpPetDChl pAmp2GChlGFP) grew more slowly than cultures containing plasmids based on minicircles with less abundant transcripts (pAmpChl and pAmpAtpBChl).

**Table 2.**
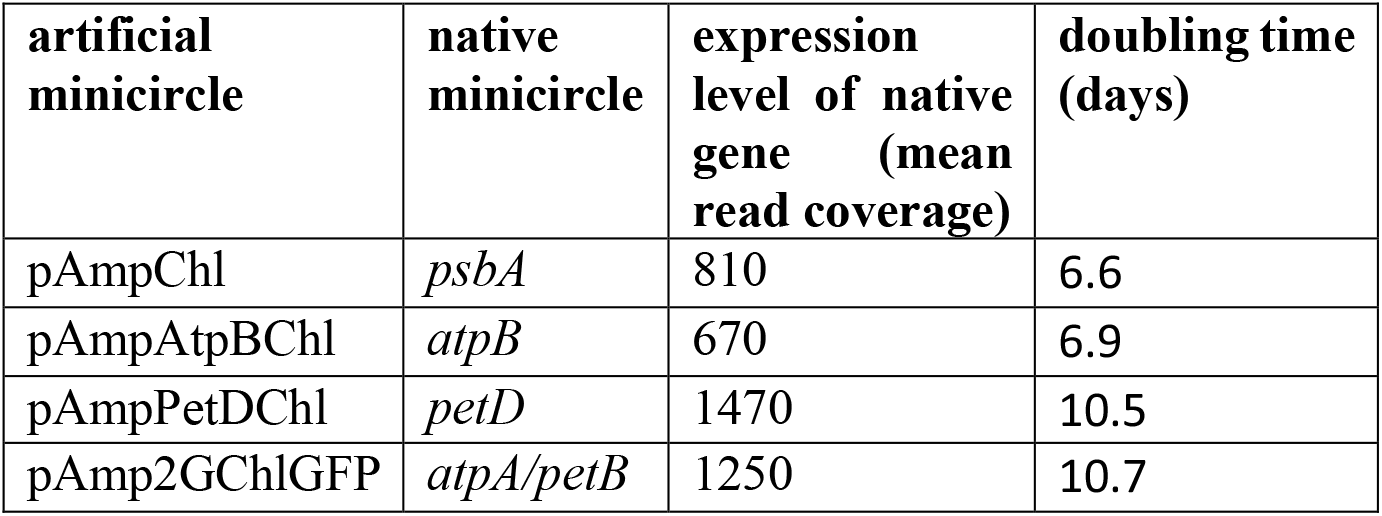
Expression of CAT in artificial minicircles. As a comparison, the levels of expression of the native minicircle from wild type cells are taken from (Dorrell et al., 2019).

| artificial minicircle | native minicircle | expression level of native gene (mean read coverage) | doubling time (days) |
| --- | --- | --- | --- |
| pAmpChl | *psbA* | 810 | 6.6 |
| pAmpAtpBChl | *atpB* | 670 | 6.9 |
| pAmpPetDChl | *petD* | 1470 | 10.5 |
| pAmp2GChlGFP | *atpA/petB* | 1250 | 10.7 |

### Expression of GFP in A. carterae containing pAmp2GChl

To determine if GFP was expressed from the two gene minicircle, pAmp2GChl, we used RT-PCR with RNA extracted from three independent cultures transformed with the plasmid and selected with chloramphenicol, as shown in Figure 4. This confirmed that GFP transcripts were present. However, it was not possible to detect functional GFP via fluorescence wide-field microscopy, due to the high levels of background fluorescence in *A. carterae* (data not shown).

**Figure 4.**
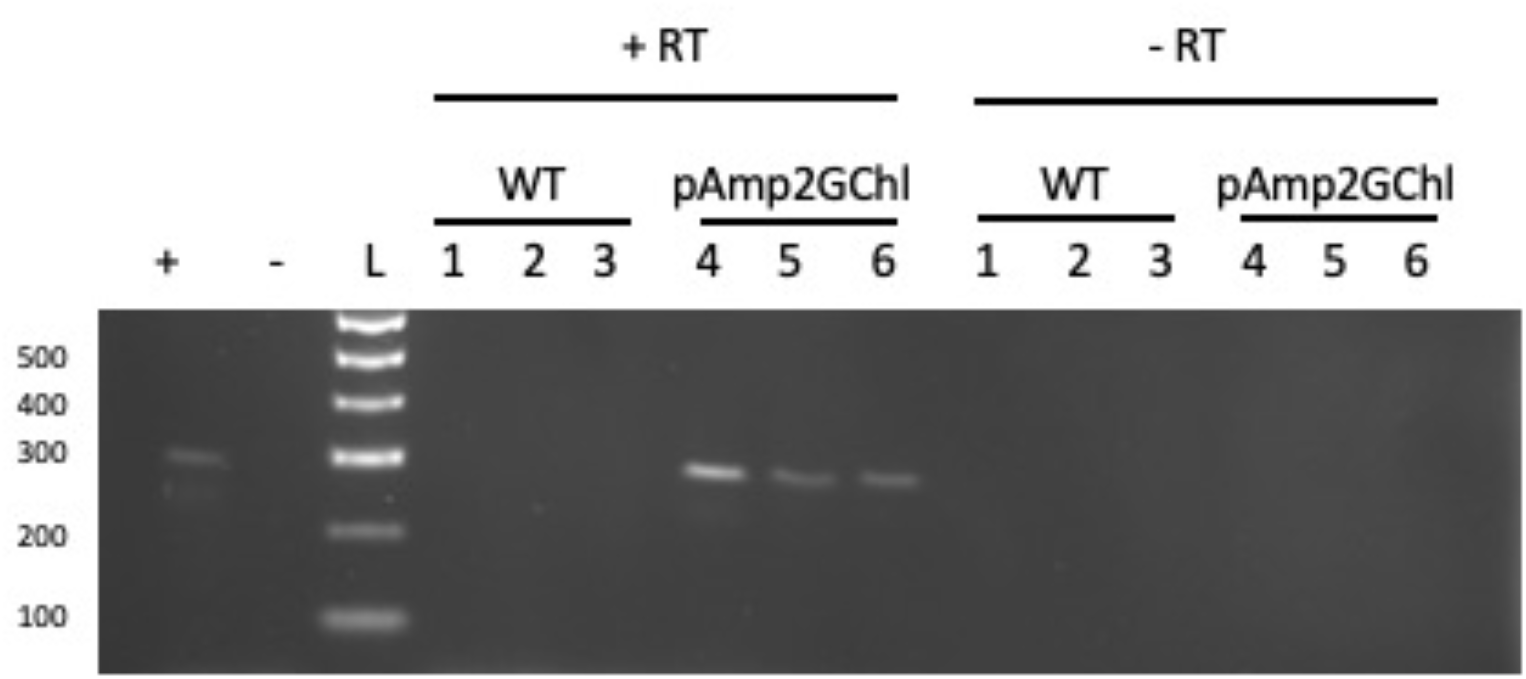
Expression of GFP in cells containing *pAmp2GChl*. RNA was extracted from wild type *A. carterae* (lanes 1-3) and cells containing pAmp2GChl (lanes 4-6). A RT-PCR reaction was carried out using primers against GFP, with and without reverse transcriptase (+RT, -RT) as shown. Lanes + and – show the results of PCR with the same primers with (+) or without (-) purified vector as DNA template.

## Discussion

The results presented here provide a significant development in the establishment of transformation technologies for dinoflagellate algae. We have doubled the number of chloroplast transformation vectors available, showing that four separate minicircles can be used for transformation. The inclusion of a two-gene artificial minicircle allows both a selectable marker and a different heterologous gene to be expressed simultaneously. However, we also find that the use of GFP as a visual marker is not suitable for use in *A. carterae*, due to the high levels of autofluorescence. These vectors will be valuable for research in a range of areas, including photosynthetic functions in dinoflagellate algae, as well determining how the fragmented chloroplast genome is maintained.

