## Supplementary material for "Establishment of a novel set of vectors for transformation of the dinoflagellate *Amphidinium carterae*": Suppl. data

**NIMMO et al Supplementary Data file**

>pAmp2GChlGFP

CTAAATTGTAAGCGTTAATATTTTGTTAAAATTCGCGTTAAATTTTTGTTAAATCAGCTCATTTTTTAACCAATAGGCCG

AAATCGGCAAAATCCCTTATAAATCAAAAGAATAGACCGAGATAGGGTTGAGTGGCCGCTACAGGGCGCTCCCATTCGCC

ATTCAGGCTGCGCAACTGTTGGGAAGGGCGTTTCGGTGCGGGCCTCTTCGCTATTACGCCAGCTGGCGAAAGGGGGATGT

GCTGCAAGGCGATTAAGTTGGGTAACGCCAGGGTTTTCCCAGTCACGACGTTGTAAAACGACGGCCAGTGAGCGCGACGT

AATACGACTCACTATAGGGCGAATTGGCGGAAGGCCGTCAAGGCCACGTGTCTTGTCCGCGGTACCTAAGTAAAGGAACT

GGAATGGTAAGGTAAATAAGGTTAAATCAAATCGAATGAAGGATACGGAAAGGTATGAAAGTCAAAACAGCGCTGAGCGT

TGACTTTCGAAGGTTAACCAACCGTAGGGATGTCAATCCCACAGGGCTCAACAACCCAGGACAAATGTTGTCAGAAGAAC

AGTTGACCTGGGCAATCATATCTGTCTCGATCATTTTGTTCCATCTCTACCCCAGTAGAGAAAAATCCAGGTCATATCAT

AGGAGATGGAACTGAGAGAACGAAGATGAAGACGAGGCGAAGAAGGATGAGGTCCCGATCTCACAAGTCTCCATTAAGGA

CTAGAGCTTGAAAGACAAATGAACGACTAGATAAATGTAGAAACGACACTTAATAGAAAGATAAGATGAAAGGCTCCCCA

TCAACACAGTACCATGAGTGCAGAGTCACTGTTGGGAGTCGAGAGATACAATATGGTCTCGAATTAATTAACAAATGCCT

CCTGGTTAGGCCAGAAGTATACCAGGAACCCTCATGCCTATTCGTTAACAAGCATTCTCTACATGGAAAAGAAGATCACC

GGTTACACAACCGTAGATATTTCTCAGTGGCATCGTAAGGAACATTTCGAGGCATTCCAGTCTGTAGCACAGTGTACATA

TAACCAGACAGTACAGCTTGATATCACCGCATTCCTTAAGACAGTAAAGAAGAACAAGCACAAGTTCTACCCAGCATTCA

TCCATATCCTTGCACGTCTTATGAACGCACATCCAGAGTTCCGTATGGCAATGAAGGATGGTGAGCTTGTCATCTGGGAT

TCTGTACATCCATGTTATACAGTATTCCATGAGCAGACAGAGACATTCTCTTCCCTTTGGTCTGAGTATCATGATGATTT

CCGTCAGTTCCTTCATATCTACTCTCAGGATGTAGCATGTTATGGTGAGAACCTTGCATATTTCCCAAAGGGTTTCATCG

AAAATATGTTCTTCGTATCTGCAAACCCTTGGGTATCTTTCACATCTTTCGATCTTAACGTCGCGAACATGGACAATTTC

TTCGCACCAGTATTCACCATGGGTAAGTATTATACACAGGGTGATAAGGTCCTTATGCCACTTGCAATCCAGGTACATCA

TGCAGTATGTGATGGTTTCCATGTAGGTCGTATGCTTAACGAGCTTCAGCAGTATTGTGACGAGTGGCAAGGTGGTGCAT

AGACTTCTATTCTATTTGAAAGGGAACTATTCTCCCGAGGCATTATTAGTGACACTAACAAGAGTACTTGTACTATCTCG

AACTACGCTGCTATGTCTAAGGGTGAAGAACTTTTCACAGGTGTAGTACCAATCCTTGTCGAGCTTGATGGTGATGTAAA

CGGTCATAAGTTCTCTGTATCTGGTGAAGGTGAGGGTGATGCAACATATGGTAAGCTCACACTTAAGTTCATCTGCACAA

CAGGTAAGCTTCCAGTACCTTGGCCAACACTTGTAACAACACTTACATATGGTGTCCAGTGTTTCTCTCGTTATCCAGAT

CATATGAAGCGTCACGATTTCTTCAAGTCTGCAATGCCAGAGGGTTATGTACAAGAGCGTACAATCTCTTTCAAGGATGA

TGGTAACTATAAGACACGTGCAGAGGTAAAGTTCGAGGGTGATACACTTGTCAACCGTATCGAGCTTAAGGGTATCGATT

TCAAGGAAGATGGTAATATCCTTGGTCACAAGCTTGAGTATAACTACAACTCCCATAACGTCTATATCACCGCAGATAAG

CAGAAGAACGGTATCAAGGCAAACTTCAAGACACGTCATAACATCGAGGATGGTGGTGTACAGCTTGCAGATCATTATCA

GCAGAACACACCAATCGGTGATGGTCCAGTACTTCTTCCAGATAACCATTATCTTTCCACACAGTCTGCACTTTCTAAGG

ATCCAAACGAGAAGCGTGATCACATGGTACTTCTCGAATTCGTAACAGCAGCAGGTATCACACATGGTATGGATGAGCTT

TATAAGTAGTTCTGTATAAAGTGTGGTTGAACGATTAGTGAGCTCGGAGCACAAGACTGGCCTCATGGGCCTTCCGCTCA

CTGCCCGCTTTCCAGTCGGGAAACCTGTCGTGCCAGCTGCATTAACATGGTCATAGCTGTTTCCTTGCGTATTGGGCGCT

CTCCGCTTCCTCGCTCACTGACTCGCTGCGCTCGGTCGTTCGGGTAAAGCCTGGGGTGCCTAATGAGCAAAAGGCCAGCA

AAAGGCCAGGAACCGTAAAAAGGCCGCGTTGCTGGCGTTTTTCCATAGGCTCCGCCCCCCTGACGAGCATCACAAAAATC

GACGCTCAAGTCAGAGGTGGCGAAACCCGACAGGACTATAAAGATACCAGGCGTTTCCCCCTGGAAGCTCCCTCGTGCGC

TCTCCTGTTCCGACCCTGCCGCTTACCGGATACCTGTCCGCCTTTCTCCCTTCGGGAAGCGTGGCGCTTTCTCATAGCTC

ACGCTGTAGGTATCTCAGTTCGGTGTAGGTCGTTCGCTCCAAGCTGGGCTGTGTGCACGAACCCCCCGTTCAGCCCGACC

GCTGCGCCTTATCCGGTAACTATCGTCTTGAGTCCAACCCGGTAAGACACGACTTATCGCCACTGGCAGCAGCCACTGGT

AACAGGATTAGCAGAGCGAGGTATGTAGGCGGTGCTACAGAGTTCTTGAAGTGGTGGCCTAACTACGGCTACACTAGAAG

AACAGTATTTGGTATCTGCGCTCTGCTGAAGCCAGTTACCTTCGGAAAAAGAGTTGGTAGCTCTTGATCCGGCAAACAAA

CCACCGCTGGTAGCGGTGGTTTTTTTGTTTGCAAGCAGCAGATTACGCGCAGAAAAAAAGGATCTCAAGAAGATCCTTTG

ATCTTTTCTACGGGGTCTGACGCTCAGTGGAACGAAAACTCACGTTAAGGGATTTTGGTCATGAGATTATCAAAAAGGAT

CTTCACCTAGATCCTTTTAAATTAAAAATGAAGTTTTAAATCAATCTAAAGTATATATGAGTAAACTTGGTCTGACAGTT

ACCAATGCTTAATCAGTGAGGCACCTATCTCAGCGATCTGTCTATTTCGTTCATCCATAGTTGCCTGACTCCCCGTCGTG

TAGATAACTACGATACGGGAGGGCTTACCATCTGGCCCCAGTGCTGCAATGATACCGCGAGAACCACGCTCACCGGCTCC

AGATTTATCAGCAATAAACCAGCCAGCCGGAAGGGCCGAGCGCAGAAGTGGTCCTGCAACTTTATCCGCCTCCATCCAGT

CTATTAATTGTTGCCGGGAAGCTAGAGTAAGTAGTTCGCCAGTTAATAGTTTGCGCAACGTTGTTGCCATTGCTACAGGC

ATCGTGGTGTCACGCTCGTCGTTTGGTATGGCTTCATTCAGCTCCGGTTCCCAACGATCAAGGCGAGTTACATGATCCCC

CATGTTGTGCAAAAAAGCGGTTAGCTCCTTCGGTCCTCCGATCGTTGTCAGAAGTAAGTTGGCCGCAGTGTTATCACTCA

TGGTTATGGCAGCACTGCATAATTCTCTTACTGTCATGCCATCCGTAAGATGCTTTTCTGTGACTGGTGAGTACTCAACC

AAGTCATTCTGAGAATAGTGTATGCGGCGACCGAGTTGCTCTTGCCCGGCGTCAATACGGGATAATACCGCGCCACATAG

CAGAACTTTAAAAGTGCTCATCATTGGAAAACGTTCTTCGGGGCGAAAACTCTCAAGGATCTTACCGCTGTTGAGATCCA

GTTCGATGTAACCCACTCGTGCACCCAACTGATCTTCAGCATCTTTTACTTTCACCAGCGTTTCTGGGTGAGCAAAAACA

GGAAGGCAAAATGCCGCAAAAAAGGGAATAAGGGCGACACGGAAATGTTGAATACTCATACTCTTCCTTTTTCAATATTA

TTGAAGCATTTATCAGGGTTATTGTCTCATGAGCGGATACATATTTGAATGTATTTAGAAAAATAAACAAATAGGGGTTC

CGCGCACATTTCCCCGAAAAGTGCCAC

| **Name** | **Type** | **Minimum** | **Maximum** | **Length** | **Direction** |
| --- | --- | --- | --- | --- | --- |
| **AmpR** | CDS | 3439 | 4299 | 861 | reverse |
| **ori** | rep_origin | 2624 | 3291 | 668 | reverse |
| **GFP** | gene | 1693 | 2409 | 717 | forward |
| **intergenic** | motif | 1603 | 1691 | 89 | forward |
| **CAT** | gene | 943 | 1602 | 660 | forward |
| **core** | motif | 488 | 708 | 221 | forward |
| **minicircle origin** | misc_feature | 387 | 2439 | 2053 | forward |

>pAmpPetDChl

CTAAATTGTAAGCGTTAATATTTTGTTAAAATTCGCGTTAAATTTTTGTTAAATCAGCTCATTTTTTAACCAATAGGCCG

AAATCGGCAAAATCCCTTATAAATCAAAAGAATAGACCGAGATAGGGTTGAGTGGCCGCTACAGGGCGCTCCCATTCGCC

ATTCAGGCTGCGCAACTGTTGGGAAGGGCGTTTCGGTGCGGGCCTCTTCGCTATTACGCCAGCTGGCGAAAGGGGGATGT

GCTGCAAGGCGATTAAGTTGGGTAACGCCAGGGTTTTCCCAGTCACGACGTTGTAAAACGACGGCCAGTGAGCGCGACGT

AATACGACTCACTATAGGGCGAATTGGCGGAAGGCCGTCAAGGCCACGTGTCTTGTCCGCAAGCTTTAATGTTGACGAAG

GATAACGAAGGCTGTTGAAGTCACAATCGACAGGAGTGATTAACTGAGGACAAATGTTGTCAGACAACCCCTCGTCCTGG

TCATGTGATAGGCTTCTCGATGAAACTGTTCCATCTCTACCCCAGTAGAGAAAAATCCAGGTCATATCATAGGAGATAGA

AATGAATGACGAGAACGAAGACAGAATGACGTTGAAGAGAACGATTAGGTAATATGTAGAAACGACAATGAAAGTCCTCT

AAAGAAGAGGCAACTAATAAAGATGAAACATTCCGACTACATGTCGAAGGCCTCTTGGTTAGGCTCTCAGACAGGTGACC

GTGCCACCGAGCATGTTATCACCTTGAAGAGATACCAGCGCTCTATCTGGTTCGGTAGCTGTTGACTCCTCTTCTCCTCT

ATTGGTTAAGGCCACGTCATCAACAAGGCTGCTTGAACTCCTCACTAGTTAATGTGGTCACTAACGGTGTTGTATTGTCA

ACCACGAGTACTGTGGCTATTAATAATGGTTATCAGCGCTATCTTCTTCTACATCTGGTTACTGTAGTGGTATCTATCTC

CTTTGACAAAGGGCCACGATACCTATGCCTCTAAGGCACCTGCAAATACCCTCTTGCGATCGAATGTGGTCAATGGGGCA

AGAGTACATGGATACCTTAATTGGTAACTAACCACGTCTATATCTGTTCGGCAGTGATTCCTATCTCCGTTCCCTTCTTG

GGATTGGGCCACAGTCTCATGCCTTAGTTGGTACTGTTTGAACCCTCATCTATCGAAAGCGGTCAATGATGATGTTGTAT

CCCTTCTGGGGCTTACCGCGGTCCACTGCATTGGCGCTGTTGATACATCCAGATGATATCTCGTCCTCTGTGTCCTATGG

GATACATGTCGTCTTCCCTCATTTGTTCATCAGTTGCCTCACCAGCTGACCTGTTACCTATGTGTCTTCGGATTAATGGA

AAAGAAGATCACCGGTTACACAACCGTAGATATTTCTCAGTGGCATCGTAAGGAACATTTCGAGGCATTCCAGTCTGTAG

CACAGTGTACATATAACCAGACAGTACAGCTTGATATCACCGCATTCCTTAAGACAGTAAAGAAGAACAAGCACAAGTTC

TACCCAGCATTCATCCATATCCTTGCACGTCTTATGAACGCACATCCAGAGTTCCGTATGGCAATGAAGGATGGTGAGCT

TGTCATCTGGGATTCTGTACATCCATGTTATACAGTATTCCATGAGCAGACAGAGACATTCTCTTCCCTTTGGTCTGAGT

ATCATGATGATTTCCGTCAGTTCCTTCATATCTACTCTCAGGATGTAGCATGTTATGGTGAGAACCTTGCATATTTCCCA

AAGGGTTTCATCGAAAATATGTTCTTCGTATCTGCAAACCCTTGGGTATCTTTCACATCTTTCGATCTTAACGTCGCGAA

CATGGACAATTTCTTCGCACCAGTATTCACCATGGGTAAGTATTATACACAGGGTGATAAGGTCCTTATGCCACTTGCAA

TCCAGGTACATCATGCAGTATGTGATGGTTTCCATGTAGGTCGTATGCTTAACGAGCTTCAGCAGTATTGTGACGAGTGG

CAAGGTGGTGCATAGTGCTACTTAACCTCCTTTACCTCAACAAGTACTTCGGTGCGAAAGGGTCTAGGAAGGTGGCTGCC

TATCTATCGCTTGGAGACGTTCCACCTATCCTCTTTCTAGAGAAGGTGATCTTGTCTACCAGGATCAGTGCGTCCCTCTT

GAACTACTCATGGAAGCAACGTAGGAGGGGATCTAATCTCACACTGTCTAACATGGCCTCGCCAGTCTTCACTAAGACCA

AATCCCTCAATAAGCTGTCTCTTTCCATCCTCAATAGGTTTAATTAATGCTGTCTCGTCTACCTCAGAGTGCGAGGGTTG

CATGGACTAGGCTCAATCGTATTAAGGTTACCTCTTCTGGTACCTTTGGTCTCAATATCAACTCGGGTTTCACCATTGGT

GCACTTCGCTTTCATCGTGATCTCTCCCTTCTCGCCCTCAAGCGATCAAAGGGTTTCACAGCCTCTAAAGCATCCGCTAA

GGCTCTTCAGCGTGCCCGTTATCGTGCCATCTCTACGTTGTCTGCCTTTCCACAGTCCATTTCCCTTGTCACGAGTTGCT

ACCACAAGCATCAGGCCTGTGTGCAGTTGTTCCCGTATATCTATCTTACTCAGGCTTGTTCTAAGTTAAGGGTTGCCTCG

AAGGTCTTGCAGACTATCTATGGCTCCGGGTTTACTGCGAGGTCTATTCTTAAGGGTTCAGCTAGTAATGCACAATTCTC

CATTCTCCTCAGTCCATAATGAAAAGGTATCGTATTTGCGTACATGGGTCGAGTGTACTCTTCCTAATGTTGGCTACTCC

CGCTGTAAAGGAGACTGCCCCTTCAACCGCAACGAGGGCAATGGTTCTTGGTGGACCAGATGTACCGGATAGTTACTTCT

TTAAGGCGAACTCATCTTTGTTTAGCCATGGTATCAACTATTCCCACAACGTCGTCTATCGTCAAGTGAAGGGACTGCTG

TCAAGGTCATCAATGCAGTAACTTAATTGGCGCGCCGGAGCACAAGACTGGCCTCATGGGCCTTCCGCTCACTGCCCGCT

TTCCAGTCGGGAAACCTGTCGTGCCAGCTGCATTAACATGGTCATAGCTGTTTCCTTGCGTATTGGGCGCTCTCCGCTTC

CTCGCTCACTGACTCGCTGCGCTCGGTCGTTCGGGTAAAGCCTGGGGTGCCTAATGAGCAAAAGGCCAGCAAAAGGCCAG

GAACCGTAAAAAGGCCGCGTTGCTGGCGTTTTTCCATAGGCTCCGCCCCCCTGACGAGCATCACAAAAATCGACGCTCAA

GTCAGAGGTGGCGAAACCCGACAGGACTATAAAGATACCAGGCGTTTCCCCCTGGAAGCTCCCTCGTGCGCTCTCCTGTT

CCGACCCTGCCGCTTACCGGATACCTGTCCGCCTTTCTCCCTTCGGGAAGCGTGGCGCTTTCTCATAGCTCACGCTGTAG

GTATCTCAGTTCGGTGTAGGTCGTTCGCTCCAAGCTGGGCTGTGTGCACGAACCCCCCGTTCAGCCCGACCGCTGCGCCT

TATCCGGTAACTATCGTCTTGAGTCCAACCCGGTAAGACACGACTTATCGCCACTGGCAGCAGCCACTGGTAACAGGATT

AGCAGAGCGAGGTATGTAGGCGGTGCTACAGAGTTCTTGAAGTGGTGGCCTAACTACGGCTACACTAGAAGAACAGTATT

TGGTATCTGCGCTCTGCTGAAGCCAGTTACCTTCGGAAAAAGAGTTGGTAGCTCTTGATCCGGCAAACAAACCACCGCTG

GTAGCGGTGGTTTTTTTGTTTGCAAGCAGCAGATTACGCGCAGAAAAAAAGGATCTCAAGAAGATCCTTTGATCTTTTCT

ACGGGGTCTGACGCTCAGTGGAACGAAAACTCACGTTAAGGGATTTTGGTCATGAGATTATCAAAAAGGATCTTCACCTA

GATCCTTTTAAATTAAAAATGAAGTTTTAAATCAATCTAAAGTATATATGAGTAAACTTGGTCTGACAGTTACCAATGCT

TAATCAGTGAGGCACCTATCTCAGCGATCTGTCTATTTCGTTCATCCATAGTTGCCTGACTCCCCGTCGTGTAGATAACT

ACGATACGGGAGGGCTTACCATCTGGCCCCAGTGCTGCAATGATACCGCGAGAACCACGCTCACCGGCTCCAGATTTATC

AGCAATAAACCAGCCAGCCGGAAGGGCCGAGCGCAGAAGTGGTCCTGCAACTTTATCCGCCTCCATCCAGTCTATTAATT

GTTGCCGGGAAGCTAGAGTAAGTAGTTCGCCAGTTAATAGTTTGCGCAACGTTGTTGCCATTGCTACAGGCATCGTGGTG

TCACGCTCGTCGTTTGGTATGGCTTCATTCAGCTCCGGTTCCCAACGATCAAGGCGAGTTACATGATCCCCCATGTTGTG

CAAAAAAGCGGTTAGCTCCTTCGGTCCTCCGATCGTTGTCAGAAGTAAGTTGGCCGCAGTGTTATCACTCATGGTTATGG

CAGCACTGCATAATTCTCTTACTGTCATGCCATCCGTAAGATGCTTTTCTGTGACTGGTGAGTACTCAACCAAGTCATTC

TGAGAATAGTGTATGCGGCGACCGAGTTGCTCTTGCCCGGCGTCAATACGGGATAATACCGCGCCACATAGCAGAACTTT

AAAAGTGCTCATCATTGGAAAACGTTCTTCGGGGCGAAAACTCTCAAGGATCTTACCGCTGTTGAGATCCAGTTCGATGT

AACCCACTCGTGCACCCAACTGATCTTCAGCATCTTTTACTTTCACCAGCGTTTCTGGGTGAGCAAAAACAGGAAGGCAA

AATGCCGCAAAAAAGGGAATAAGGGCGACACGGAAATGTTGAATACTCATACTCTTCCTTTTTCAATATTATTGAAGCAT

TTATCAGGGTTATTGTCTCATGAGCGGATACATATTTGAATGTATTTAGAAAAATAAACAAATAGGGGTTCCGCGCACAT

TTCCCCGAAAAGTGCCAC

| **Name** | **Type** | **Minimum** | **Maximum** | **Length** | **Direction** |
| --- | --- | --- | --- | --- | --- |
| **AmpR** | CDS | 3990 | 4850 | 861 | reverse |
| **ColE1origin** | rep_origin | 3175 | 3842 | 668 | reverse |
| **CAT CDS** | misc_feature | 1356 | 2015 | 660 | forward |
| **minicircle backbone** | CDS | 387 | 2988 | 2602 | forward |
